# Echtvar: Compressed variant representation for rapid annotation and filtering of SNPs and indels

**DOI:** 10.1101/2022.04.15.488439

**Authors:** Brent S. Pedersen, Jeroen de Ridder

**Author notes:** Correspondence &.

## Abstract

Germline and somatic variants within an individual or cohort are interpreted with information from large cohorts. Annotation with this information becomes a computational bottleneck as population sets grow to terabytes of data. Here, we introduce *echtvar*, which efficiently encodes population variants and annotation fields into a compressed archive that can be used for rapid variant annotation and filtering. Most variants, including position and alleles are encoded into 32-bits–half the size of previous encoding schemes and at least 4 times smaller than a naive encoding. The annotations, stored separately, are also encoded and compressed. We show that echtvar is faster and uses less space than existing tools and that it can effectively reduce the number of candidate variants. We give examples on germ-line and somatic variants to document how *echtvar* can facilitate exploratory data analysis on genetic variants. *Echtvar* is available at https://github.com/brentp/echtvar under an MIT license.

## Introduction

A site in the genome that differs from the reference, either as a somatic mutation or a germline variant must be decorated with additional information in order to be interpretable. Millions of sites in an individual will differ from the reference genome. Several pieces of information can be added to each variant to assist in determining which of those are relevant to disease. It is critical to know the predicted effect on a gene–for example, does it create a new stop-codon in the sequence of an exon? Additionally, the frequency or absence of a variant in a large population database indicates potential constraint with the species^1^. Likewise, the conservation of the site across species^2^ indicates that a site might be important and so those should experience selection and be removed from the population. Each of these pieces of information must be added to each variant in a call-set using an annotation tool.

Tools that *calculate* the effect of a genetic variant on protein (and non-coding) sequence such as Variant-effect predictor (VEP)^3^, bcftools csq^4^, and snpEff^5^ are invaluable; but here, we focus on the annotation that involves *searching* for a particular variant or site in a database and annotating the variant with this information from the match in the database. This way of annotating variants is a fundamental building block in most genetics data analysis pipelines and plays a critical role in variant interpretation. For example it is very common to annotate with population allele frequency from the Genome Aggregation Database (gnomAD)^6^ or other large population sets. Another example is the addition of a CADD^7^ score for each variant in a Variant Call Format (VCF) file^8^. While conceptually simple, the space and time efficiency of the annotation algorithms become critical as call-sets and annotation databases have grown substantially. The methods involved in annotating with the essential data in these large datasets become quite relevant. As the size of the annotation sets grow, the space and time efficiency of the algorithms become more important. The annotations added by the tools that implement these algorithms are fundamental to variant interpretation.

As an example of the scale of the data, the Genome Aggregation Database (GnomAD)^6^ v3.1.2, mentioned above, contains nearly 760 million variants, consuming more than 2 terabytes of data. Storing this database can be onerous on an average compute cluster and hard to justify when the size of alignment and variant information for a trio that an investigator might wish to annotate is on the order of a few gigabytes of data. Further, attaching population information to each variant in this example trio would take additional compute to decompress and parse the huge gnomAD files, even when using an approach that combines index-jumping and streaming like VCFAnno^9^. Likewise, the CADD^7^ score includes a prediction for each of 3 possible single-base changes for each position in the human reference; this commonly-used annotation is 81 Gigabytes of compressed data and incurs substantial compute. Even annotations for only coding variants can be quite large, for example dbNSFP^10^, which aggregates many of these scores, totals around 30 Gigabytes of compressed data. Data this size require new methods in order to be utilized with efficiency and ease, especially given their routine use in modern day genetics pipelines.

Here, we introduce our command-line tool, *echtvar*, and show how the annotations and filtering performed by *echtvar* can dramatically reduce the number of candidate variants and facilitate exploratory data analysis due to the speed and ease of use. We document performance in terms of speed, memory, and number of variants filtered, in experiments looking at germline and somatic variants. *Echtvar* can ease the interpretation of whole-genome sequencing by quickly limiting the number of variants under consideration for both germ-line and somatic projects. This reduces the cost of computation and downstream analysis. Such advances can facilitate efforts to improve turn-around times for sequencing projects where speed is critical, such as for neonatal projects^11^.

### Results

We first give a brief summary of the *echtvar* algorithm (this is expanded in the methods section), then we compare *echtvar* to other tools on a practical example–annotating a set of whole-genome germline variants with information from gnomAD. This demonstrates the speed and memory use of *echtvar* relative to other tools on a common, yet sizable, task. Then, on the same germline variants we show the filtering capabilities of *echtvar* which enable interactive, exploratory data-analysis. Finally, we give an example of using *echtvar* to annotate somatic variants of thousands of samples from International Cancer Genome Consortium (ICGC) with values from dbNSFP^10^.

### Brief algorithm overview

*Echtvar* chunks the genome, efficiently encodes variants into integers, and utilizes integer compression methods to facilitate compact variant representations that can be used for rapid annotation. Briefly, *echtvar* combines the following:

1. an encoding scheme that fits most variants, including position, reference, and alternate alleles into 32 bit integers
2. a chromosome and region chunking for file-layout scheme that limits memory-use and improves compression
3. the Stream-VByte encoding scheme^12^ which can encode and decode billions of integers per second while reducing the space required by nearly 4 times for some field-types
4. use of the standard ZIP file format to allow random-access to each region
5. a command-line interface that allows users to create custom *echtvar* archives by extracting specific integer, float, and low-arity string fields from population databases.
6. a tool to annotate and filter query variant files with values in *echtvar* archives.

This combination of methods and utilities make *echtvar* a valuable tool for annotating and filtering genetic variants. We expand on the process of encoding and annotation in more detail in the methods.

### Comparison with other tools

We compare *echtvar* speed, memory-use and archive size to bcftools annotate ^4^, VarNote^13^, and slivar^14^ on gnomAD v3.1.2^6^ annotating Genome-in-a-Bottle calls for HG001^15^ which contains about 3.9 million SNP and indel whole-genome, germline calls.

*Echtvar* is the fastest tool (Fig 1a, b) with the smallest annotation file-size footprint (Fig 1d) while using a small amount of memory (Fig 1c) for any modern server. *Echtvar* completes the task in 132.2 seconds with 68.1 Megabytes of memory; the closest competitor is BCFtools which uses 396.7 seconds and 43.5 Megabytes of memory. Note that *echtvar* uses only 7.3 Gigabytes on disk while BCFtools uses 12.6 Gigabytes. These sizes are close because we subset the gnomAD VCF to contain only fields of interest to make the comparison as fair as possible–the original VCF files are around 2 terabytes of data. The *echtvar* command used for this comparison was:

~~~
echtvar anno \
 -e gnomad.v3.1.2.echtvar.v2.zip \
 $vcf $output_vcf
~~~

where “$vcf” and “$output_vcf” are placeholders for the input VCF to be annotated and the output file where results are written.

**Figure 1.**
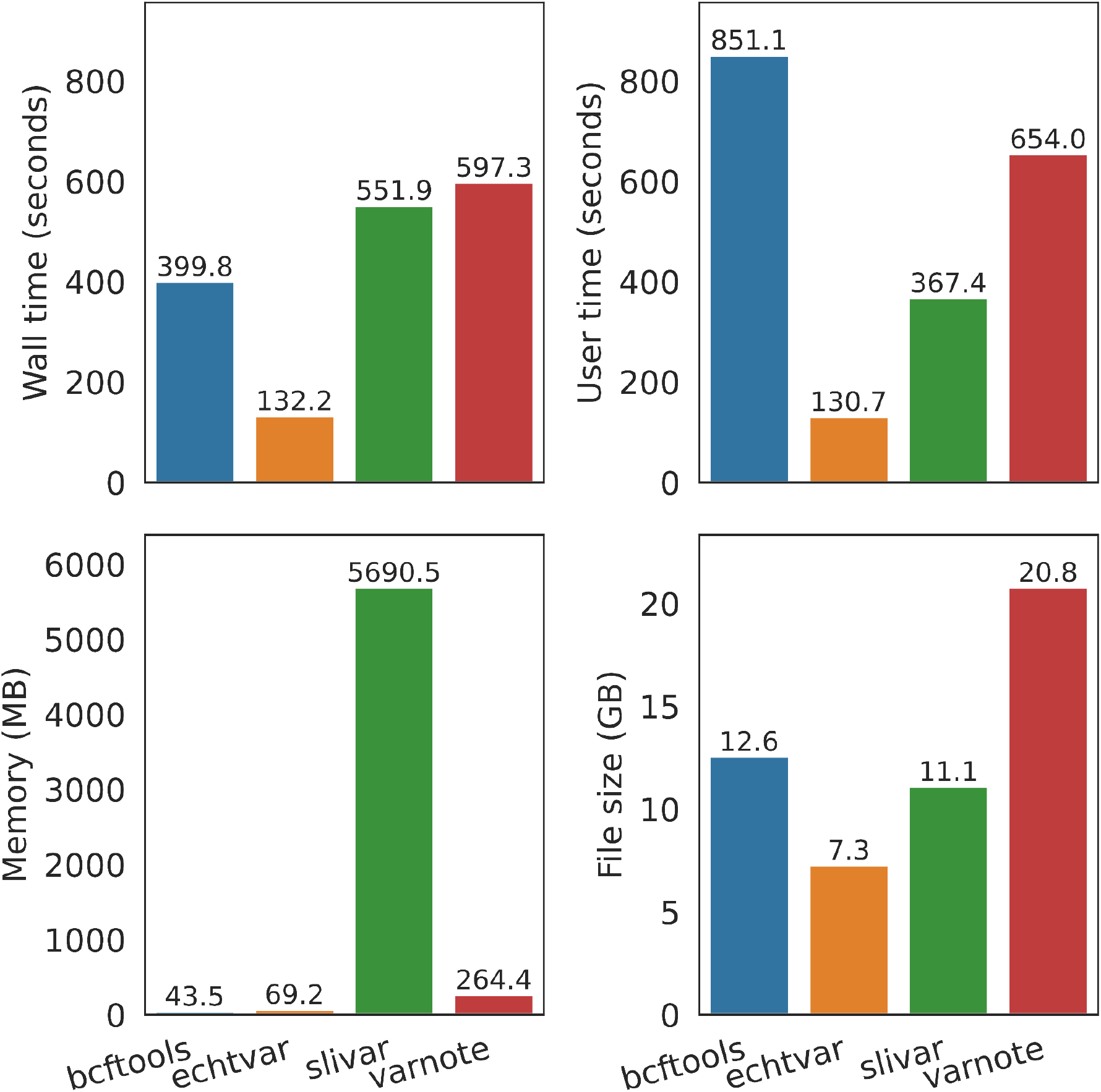
Comparison of *echtvar* speed, memory, and archive size with other annotation tools. A VCF with about 3.9 million variants was annotated with allele frequency, number of homozygous alternate alleles, and other fields from gnomAD v3.1.2. The first row shows wall-time, how long the programs took to complete (a) and user-time, how much processing time (across multiple threads) was used (b). The amount of memory used is shown in c. with *slivar* taking much more memory than other tools. d) shows the file size of the annotation files required. Since *echtvar* encodes the data, it can use less file size than the original file. Original gnomAD file sizes are much larger, the values shown for *bcftools* and *varnote* are from a VCF or BCF subset to contain only the fields of interest for a more fair comparison.

### Filtering whole-genome germline variants

Next, we remove variants unless the highest allele frequency of any population in gnomAD v3.1.2 was less than 0.01. For this purpose, we used the same set of calls for HG001 along with the same *echtvar* annotations from the previous comparison. This can be achieved with the following *echtvar* command:

~~~
echtvar anno -e gnomad.v3.1.2.echtvar.v2.zip $vcf $output_vcf \
 -i “gnomad_popmax_af < 0.01”
~~~

While exact filtering strategies will vary, this is a reasonable starting filter for rare-disease variants, where we expect candidate variants contributing to a severe phenotype to be rare. In doing this filtering, we reduced the *echtvar* run-time from 132 seconds (as in Figure 1) to 87 seconds (34%) and reduced the number of variants from around 3.95 million to 67,017 (98% reduction). The speed improvement is because fewer variants are written and writing to file is otherwise a bottleneck in the annotation step. The filtered variant set is 50 times smaller and so will potentially use 50 times less storage depending on the compression (4.2MB vs 127MB == 30 times for this example), and less compute for intensive downstream tasks such as effect annotation, for example with Variant Effect Predictor (VEP)^3^. Combining the annotation with filtering compounds the benefit of each of these steps and also highlights the utility of fast tools that leverage large population datasets such as gnomAD in prioritizing variants.

### Filtering recessive whole-genome germline variants

Next, in order to further show the capabilities of *echtvar*, we evaluate filtering for recessive variants where we expect that sites (variants) contributing to disease would have few samples from gnomAD that were homozygous for the variant. We therefore filtered to variants where the proportion of homozygous alternate samples across all populations was less than 0.5% of the number of total samples in that population. *Echtvar* supports this through the following command:

~~~
echtvar anno -e gnomad.v3.1.2.echtvar.v2.zip $vcf $output_vcf \
 -i “gnomad_popmax_nhomalt / (gnomad_popmax_an / 2) < 0.005”
~~~

Note that we get the number of samples using the number of chromosomes (the “an” suffix is for “number of alleles” across the population) divided by 2 since we are considering only the autosomes. The left-hand side then gives the proportion of samples and we compare that to the right-hand size (0.005). This completes in around 90 seconds and writes 178,117 variants (95.5% of variants filtered). This example demonstrates the flexibility of the expressions which allow a variety of mathematical operations. It also highlights the advantage of such a fast tool. We can rapidly evaluate expressions of 4 million variants to decide on the exact filtering parameters. For example if the analyst were to decide that 178 thousand variants is too many, they could run again with a cutoff of 0.1% and have the results in about a minute and a half.

### Filtering whole-genome somatic variants

To demonstrate applicability in a somatic variant setting, we annotated and filtered each of 1,902 VCF files of somatic variants from the International Cancer Genome Consortium (ICGC) with annotations from dbNSFP. We used the command:

~~~
echtvar anno -e dbNSFP.echtvar.zip $vcf /dev/null \
   -i ‘dbsnfp_SIFT_converted_rankscore > 0.2 \
   || dbsnfp_DANN_rankscore > 0.2 \
   || dbsnfp_GERPpp_RS_rankscore > 0.2 ‘
~~~

to annotate with dbNSFP and filter to variants that had at least one of the rank scores greater than 0.2. We chose these fields and expressions to highlight the flexibility and possibilities of *echtvar* rather than to address a specific question. Figure 2 shows the time required to run this command for each file, along with the number of variants left after filtering. All commands finish in a few seconds and leave only a handful of variants in most cases. This demonstrates how one could rapidly evaluate different cutoffs to get to a reasonable number of variants of interest. While nearly all samples had fewer than 50 variants that passed the filters, a few samples had more than 100 variants (not shown in Fig 2b which is truncated at 100). These could be samples that require further quality control. As *echtvar* readily achieves these calculations in a matter of seconds it would, for instance, be possible to include them as broad quality control measures that require little extra compute.

**Figure 2.**
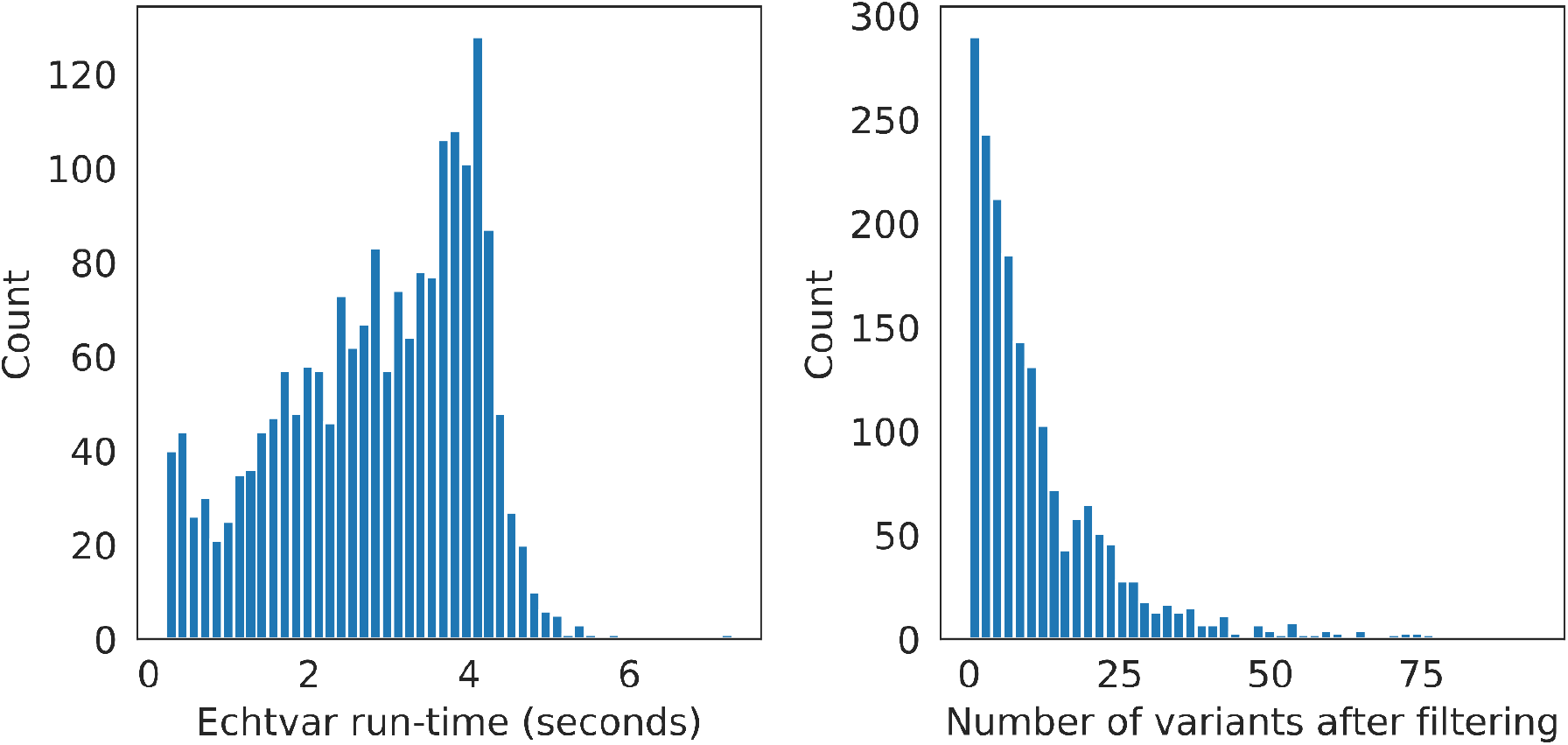
Histograms of run-time (A) and variants remaining after filtering (B) for 1,902 VCFs containing somatic variants for a variety of cancer types from the ICGC. We annotated each VCF with data from dbNSFP version 4.3a and then filtered to variants with a high score in any of 3 of the annotated fields.

### Discussion and Future Work

Variant annotations from large population sets are essential for virtually all variant interpretation and downstream analyses. We have introduced *echtvar* which uses genome-chunking and an encoding scheme that fits most SNP and indel variants into 32-bit integers to facilitate rapid and flexible annotation. We have shown that *echtvar* is at least 3 times as fast as competing tools while using less space for the encoded data and very little memory. We also showed examples of using *echtvar* to simultaneously annotate and filter germline variants; first to those with low allele frequency in a population and then to those with a small percentage of homozygous samples. These are example analyses that are common in rare-disease research. We also showed how *echtvar* can be used to filter somatic variants. In all of these, the speed and simplicity of *echtvar* make it a valuable tool for variant annotation and filtering and for exploratory data analysis.

While we did explore different encoding schemas, future work could evaluate using 64-bit integers instead of 32-bit–this would allow more variants to fit in the concise scheme at the cost of a larger average size. This increase in size could be mitigated by compression, but the delta-encoding, where each value is stored as the difference to the previous value, is less effective when fewer bits are used for compression. For example, if 20 bits are used for position, then 2 adjacent variants would differ by at least 44 bits, limiting the benefit of both delta encoding and VByte compression. Other work could explore the trade-off in using bins with a fixed number of variables, rather than a fixed size. This would mean that an index for the starting position of each bin would need to be maintained but that each bin would have a similar size in memory; this could improve changes to *echtvar* that focus on parallelization which is another area for future research.

We expect that the simplicity, speed, and utility of *echtvar* will make it a staple in variant annotation pipelines.

*Echtvar* is available under the liberal MIT license from https://github.com/brentp/echtvar. There is a static binary that will work immediately on modern linux systems.

## Methods

### *Echtvar* Encoding

*Echtvar* accepts a VCF (or BCF)^8^ and a JSON configuration file that indicates which fields should be extracted from the INFO field of each variant and how they are stored in the *echtvar* ZIP archive. This archive partitions each chromosome into 1,048,576 (2^20^) base intervals (bins) which are stored in separate directories (Figure 3). The amount of data in memory for encoding and annotation is determined by the number of variants and fields within each bin. Each bin contains one 32-bit entry for each variant from the VCF. Small variants, those with a combined reference and alternate allele length of fewer than 5 bases, are encoded into 32 bits and stored directly in the primary table (Figure 3). Because each chromosome and 1,048,576 (2^20^) base interval is stored in a separate directory within the ZIP archive, only 20 bits are needed to indicate the position of the variant within that interval (Figure 3, upper right). The remaining 12 bits in a 32 bit integer can store the reference and alternate alleles of variants with a total length (REF + ALT) of fewer than 5 bases (Figure 3, upper-right). This is possible, because with 4 total nucleotides, we only need 2 bits to store each nucleotide, but we also need to store, within those 12 bits, the length of the reference and alternate alleles. Around 92% of variants in gnomAD^6^ v3.1.2 fit into 32 bits. This encoding is similar to VariantKey^16^ but VariantKey encodes the full position along with the chromosome into a 64-bit integer.

**Figure 3.**
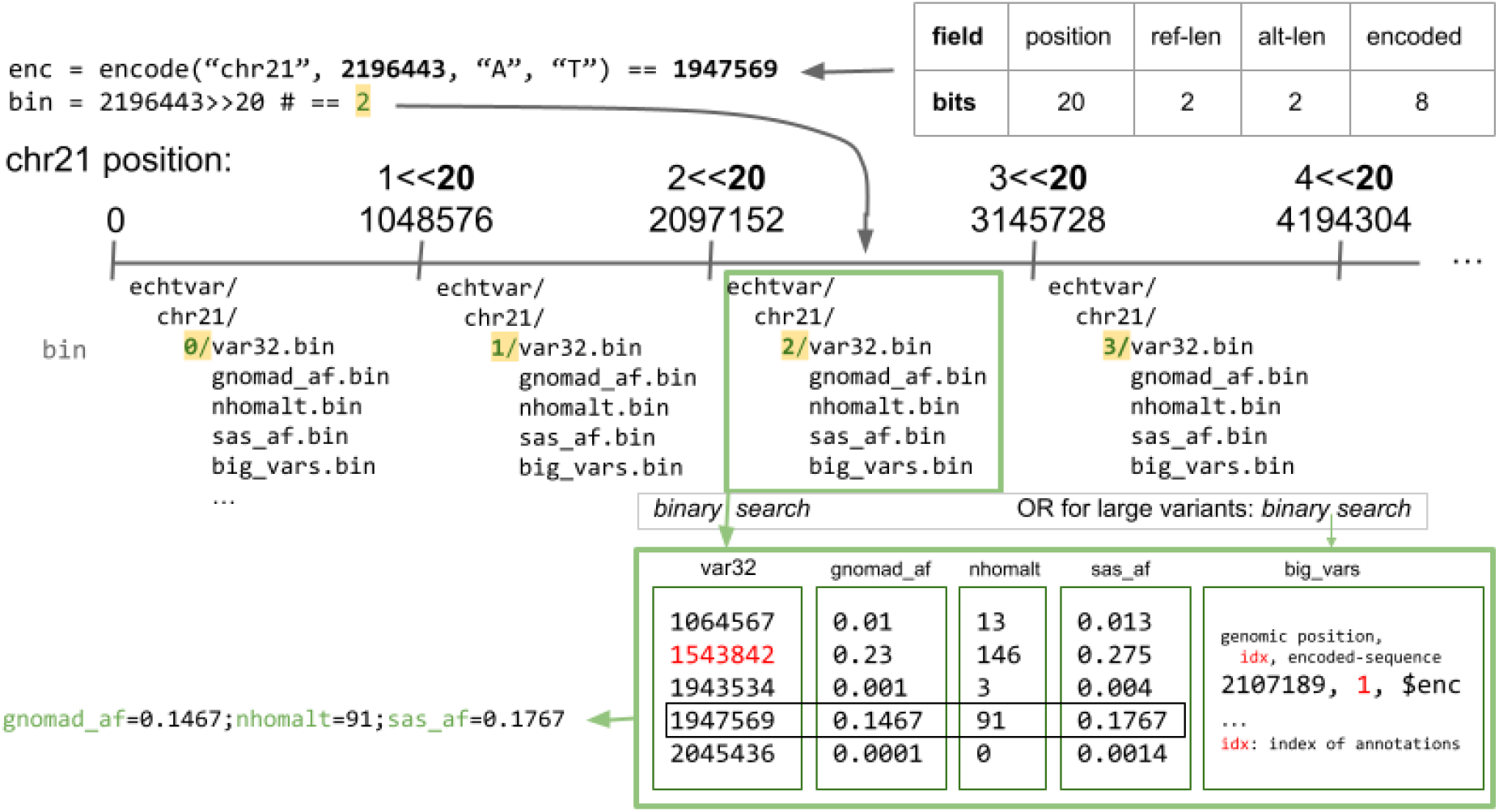
*Echtvar* encoding and annotation schema. *Echtvar* encodes small variants into 32-bit integers with the bits partitioned as in the top-right. Encoding simply partitions values to those bits which results in a 32-bit integer. The genomic bin determines the 1,048,576 bin and corresponding directory within the *echtvar* archive for a given query variant. During annotation, if the bin is different from the previous query variant, the data for that bin, including the primary (*var32*) table, the larger variants in the supplemental (*big_vars*) table, and the fields, are all read into memory. If the bin is the same, the values are already in memory. The encoded variant is then used in a binary search against the primary (*var32*) table to find the index of the variant. That index is then used to extract the corresponding fields. If the variant is not found in the table, user-specified default values are returned. Variants with a combined reference and alternate allele length greater than 4 bases will not fit into 32-bits and must be encoded and then searched in the large-variants, supplemental (*big_vars*) table. The binary search in that table again yields an index which is used to extract the associated fields. Those fields are then added to the query variant which is written to the output.

For each long variant, a place-holder variant with empty reference and alternate allele is encoded and inserted into the primary variant file for that bin. In addition, variants that are 4 bases or larger (longer insertions or deletions) are stored in a different file within each bin in a still efficient format that uses a variable-length encoding to handle any size variant.

Each field that is requested by the user, for example “AF” for allele-frequency, is extracted and encoded into a file (for each bin) with the user-specified alias, such as “gnomAD_AF”. Within each of those value files, there is one value for each variant from the VCF that falls within that region. The configuration file can also specify a default value when that file is missing, and other modifications to default parameters. Upon encoding, the user-specified configuration file is stored in the ZIP archive.

Within each bin, the encoded variants (and place-holder variants) are sorted to allow for fast searching. The variants are delta-encoded – so that only the difference between each 32 bit encoded variant and the one that precedes is stored. This requires the extra step of performing the cumulative (prefix) sum upon annotation, but improves compression. The delta-encoded variants are then further encoded with Stream VByte-encoding^12^ which encodes integers to use between 1 and 4 bytes depending on the size of the value; a separate block of “control bytes” indicates the number of bytes consumed by each integer. Since it is common to have many small numbers, especially in variant annotation where most variants are rare, this can effectively compress the data. In addition, the schema, where the control bytes are stored separately, allows modern processors to rapidly decode the data and allows skipping numbers without decoding them. Longer variants that do not fit within the 32 bits are encoded with bincode^17^ which internally uses compression and *varint* encoding. *Varint* encoding is similar to Stream VByte-encoding, except that the control bytes are stored with each encoded number.

As each variant is encoded and inserted, the corresponding user-requested INFO fields are inserted into vectors such that there is a one-to-many correspondence between a variant and the fields. Each field vector will have exactly the same number of entries as the encoded variant vector. During iteration of the VCF, once a new 2^20^ base bin is reached, the previous bin, including primary encoded variants, long variants, and all fields are written to separate files within the same ZIP directory for that bin. Fields also undergo Stream VByte-encoding; floating point values are first converted to integers by multiplying by a user-specified value. Upon annotation, the extracted integers are then divided by that same multiplier to regain nearly the same value. Higher multipliers give better precision but less compression. Integer values do not need to undergo this transformation but are limited to 32 bits.

*Echtvar* encodes string fields from a VCF into integers in the archive by using an extra lookup vector of the unique observed strings. For each unique string observed in any bin, *echtvar* inserts that string into the vector and stores the index of that vector for that variant. For low cardinality fields, for example, with only 10 unique values, this means that only integer values between 0 and 9 are saved in the field arrays. Once encoding of the entire file is complete, the string arrays are written to a single file per field. During later annotation, the string arrays are then used to convert from the integer stored per variant to the actual string value.

### Echtvar Annotation

To annotate a query VCF with an *echtvar* archive, the user specifies those two files along with an output path to write the annotated VCF (or BCF) file. All fields from the archive are added to the output file. For each variant in the query VCF, if the position is in a different bin than the previous variant, then the files for the new bin, including variants, long variants, and fields, are read into memory. As such, *echtvar* is fastest on files that are sorted by genomic position. This sorting is a requirement for the other tools we compare to. If the variant has a total length fewer than 5 bases, *echtvar* encodes the variant into a 32-bit integer and does a binary search against the primary variant table to find the index. Note that variants from the archive remain as integers and do not need to be decoded back into variants. The index from the binary search is then used to extract the values for each field and add them to the query variant (see Figure 3). Query variants with 5 or more total bases are encoded into the longer format and a binary search against the supplemental table (of long variants) is performed. That yields an object that contains an index which is then used to extract the value for each field in the archive. At this point, the extracted fields are then tested against a user-specified filter if one was given. If the filter passes (evaluates to true), then the fields are added to the query variant which is then written to the output file. The filter is evaluated using *fasteval*^*18*^.

### Libraries used in *echtvar*

These methods are achieved with the help of a number of libraries. We use HTSLib^19^ via rust-htslib^20^ to read, update and write the VCF files. We use fasteval^18^ to parse and evaluate the filter expressions, stream-vbyte-rust^21^ to perform the stream-vbyte compression, bincode^17^ to compress large variants, and zip-rs^22^ to create the *echtvar* zip archive.

### Whole-Genome Variants Annotated with gnomAD: comparison with other tools

We downloaded gnomAD v3.1.2. In order to make the comparison as fair as possible, we subset the files to contain only the 10 INFO fields of interest, and concatenate them into a single 20GB file. We used this to annotate variants from Genome in a Bottle (GIAB) for HG001 from: https://ftp-trace.ncbi.nlm.nih.gov/giab/ftp/release/NA12878_HG001/NISTv4.2.1/GRCh38/HG001_GRCh38_1_22_v4.2.1_benchmark.vcf.gz

We used bcftools^4^ norm to decompose and normalize the variants to a consistent representation.

All tools were added to a single docker image for reproducibility and versioning. For *slivar, echtvar*, and *varnote* we performed the necessary encoding steps documented in the script linked below. Since these encodings are one-time costs, we did not compare the run times. We then evaluated the tools using the commands in: https://github.com/brentp/echtvar/blob/main/paper/echtvar-paper.sh

We saved the times using /usr/bin/time -v and we also saved the total size of all files needed for the annotation.

### Filtering whole-genome variants

We used the gnomAD v3.1.2 archive described above and the HG001 query VCF to evaluate the effect of filtering. We simply added the parameter:

~~~
 -i ‘gnomad_popmax_af < 0.01’
~~~

to include only variants that met that expression. The *gnomad_popmax_af* filter is from the *AF_popmax* field in the original gnomAD VCFs that indicates the maximum allele frequency across each of the sub-populations in gnomAD. A variant contributing to a severe phenotype should be rare in all populations; using the maximum across populations allows us to apply that filter.

### Filtering somatic variants with dbNSFP

We annotated 1,902 VCF files of somatic variants with dbNSFP version 4.3a. First, we converted dbNSFP to VCF format using this script from the echtvar repo: https://github.com/brentp/echtvar/blob/main/scripts/dbnsfp.py

We then converted the resulting VCF to and echtvar archive with the following command:

~~~
echtvar encode dbNSFP.echtvar.zip dbNSFP.json $dbnsfp.vcf.gz
~~~

where dbNSFP.json contains:

~~~
[ {
  “field”: “SIFT_converted_rankscore”,
  “alias”: “dbsnfp_SIFT_converted_rankscore”,
  “multiplier”: 1000000
}, {
  “field”: “DANN_rankscore”,
  “alias”: “dbsnfp_DANN_rankscore”,
  “multiplier”: 1000000
}, {
  “field”: “GERP++_RS_rankscore”,
  “alias”: “dbsnfp_GERPpp_RS_rankscore”,
  “multiplier”: 1000000
}]
~~~

Finally we annotated each ICGC VCF with the archive using:

~~~
echtvar anno -e dbNSFP.echtvar.zip $vcf /dev/null \
  -i ‘dbsnfp_SIFT_converted_rankscore > 0.2 \
  || dbsnfp_DANN_rankscore > 0.2 \
  || dbsnfp_GERPpp_RS_rankscore > 0.2 ‘
~~~

while saving the run-time. Full commands for this are in this script: https://github.com/brentp/echtvar/blob/main/paper/icgc.sh

## Acknowledgements

We benefited from feedback from Sascha Brunner, Myrthe Jager, Arne van Hoeck, Fran Martinez and Fritz Sedlazeck.

